# The biochemical resolving power of fluorescence lifetime imaging: untangling the roles of the instrument response function and photon-statistics

**DOI:** 10.1101/2021.02.03.429635

**Authors:** Andrew L. Trinh, Alessandro Esposito

## Abstract

A deeper understanding of spatial resolution has led to innovations in microscopy and the disruption of biomedical research, as with super-resolution microscopy. To foster similar advances in time-resolved and spectral imaging, we have previously introduced the concept of ‘biochemical resolving power’ in fluorescence microscopy. Here, we apply those concepts to investigate how the instrument response function (IRF), sampling conditions, and photon-statistics limit the biochemical resolution of fluorescence lifetime microscopy. Using Fisher information analysis and Monte Carlo simulations, we reveal the complex dependencies between photon-statistics and the IRF, permitting us to quantify resolution limits that have been poorly understood (*e.g*., the minimum resolvable decay time for a given width of the IRF and photon-statistics) or previously underappreciated (*e.g*., optimization of the IRF for biochemical detection). With this work, we unravel common misunderstandings on the role of the IRF and provide theoretical insights with significant practical implications on the design and use of time-resolved instrumentation.

## 1. Introduction

Fluorescence lifetime imaging microscopy (FLIM) is an essential tool that can quantitate biochemistry in living cells with single-cell or sub-cellular resolution. The fluorescence lifetime of a fluorophore, *i.e*. the average time that a fluorophore spends in its excited state after absorbing a photon, depends on the physico-chemical environment of the fluorophore [1, 2]. The fluorescence lifetime of some fluorescent molecules is intrinsically sensitive to parameters such as pH, viscosity, and oxygen concentration. Other fluorophores are engineered to sense biologically-relevant quantities such as analyte concentrations, protein-protein interactions, conformational changes, and post-translational modifications, often measuring variations in fluorescence lifetimes mediated by Foerster Resonance Energy Transfer (FRET) [3, 4].

Time-correlated single-photon counting (TCSPC) is often regarded as the gold-standard in time-resolved detection. TCSPC measures the arrival time of individual photons relative to a train of identical excitation pulses of light. Single-photon counting techniques provide the significant advantage of being ‘shot-noise’ limited, *i.e*. their noise performance can reach the limits imposed by the quantised nature of light. The signal-to-noise ratio (SNR) in fluorescence imaging cannot exceed the limit imposed by Poissonian noise for the number of collected photons, *N* [5, 6]:

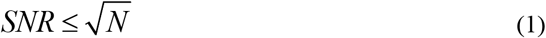

TCSPC approaches this limit albeit with losses determined by pulse pile-up effects, *i.e*. the inability of resolving a second photon while the detection system is still busy handling the electrical trace of the first (due to the combined effect of the detector and the electronics dead-times), or difficulties in detecting multiple photons per light pulse. We have made significant progress in ameliorating pulse pile-up effects, for example, by utilizing multi-hit time-to-digital [7, 8] or multiplexed time-to-analog converters [9], ultra-fast digitizers [10], fast detectors (*e.g*. hybrid photomultipliers [11]), and detector arrays [12–15]. Such innovations permit contemporary FLIM systems to acquire images with fluorescence lifetime contrast at spatial resolutions typical of confocal microscopy without significantly compromising imaging speed.

The need to reach higher acquisition speeds may require a trade-off in other instrument specifications such as the number of bins used to sample fluorescence decays or the instrument response function (IRF), *i.e*. the uncertainty over the sampled arrival time of each photon caused by variabilities in the excitation pulse, the transit time spread of the detector, and the time-jitter of the detector and electronics [16]. The IRF can be measured using samples with well-characterised fluorescence lifetime values or samples with nominal zero-lifetimes (*e.g*., scattering of the excitation light or second-harmonic generation). The measured signal, *M(t)*, is thus represented by the convolution integral of the fluorescence decay, *D(t)*, and the instrument response function, *IRF(t) [1, 2, 17, 18]*.

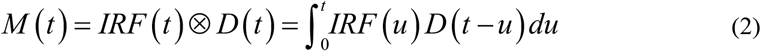

It is generally accepted that the width of the IRF limits the shortest measurable lifetime value. However, the impact of the IRF on the overall precision of fluorescence lifetime estimations is not well-characterised. The effects of other imaging parameters on the precision of TCSPC is much better understood. For example, Köllner and Wolfrum applied Fisher information theory to derive an analytical solution for the precision of a system with a given number of time channels, measurement period, and sample lifetime [5, 6, 19, 20]. Similarly, Draaijer *et al*. introduced the figure of merit, *F*, as **Eq. (3)**, to optimise the performance of time-resolved detection by adapting the position, width and number of time-bins used for histogramming the fluorescence decay [21]:

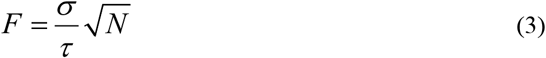

*F*, also referred to as the photon-economy, is the ratio of the relative error of the lifetime estimate (*σ* / *τ*) to the relative error due to Poissonian noise (*N*^½^ / *N* = *N*^-½^). If the lifetime estimate is only affected by Poissonian noise, then F is equal to 1, and we refer to the system as being efficient with its collected photons. In this case, the identity in **Eq. (1)** holds, and FLIM could attain the maximum SNR. The analysis of F-values has been applied to compare various implementations of FLIM [5, 23] and to optimize imaging parameter [22, 24–26].

Currently, our understanding of the resolution limit of FLIM is curtailed by our lack of a theoretical foundation. While it is generally accepted that broader IRFs result in worse resolution, a theoretical framework that describes the interdependence between the IRF width, sampling conditions, and photon-statistics is lacking. Here, we provide the Fisher information analysis and Monte Carlo simulations to fill this gap in knowledge. Given that the recent developments of fast and efficient FLIM systems within the biomedical community have been stimulated by methodological advances, such as hybrid photomultiplier tubes [11], multi-hit time-to-digital converters [7, 8], integrated smart-pixel technologies [27, 28], and ultra-fast digitizers [10], we specifically focused on how limitations in the instrument temporal resolution impacts the ‘biochemical resolving power’ [24]. We envisage that a deeper understanding of the trade-offs required for the development of FLIM systems designed for biomedical applications will aid the community to break new ground in the development and application of biochemical imaging.

## 2. Methods

### Simulations

We implemented Monte Carlo simulations in Python (www.python.org) using NumPy [29] and SciPy [30], and generated plots by Matplotlib [31]. The Jupyter notebook used to generate all the figures of this work are available at the GitHub repository *Esposito-Lab/BiochemicalResolution* [32]. We simulated fluorescence decays by generating the probability density function (PDF) of the exponential decay with lifetimes between 0.2 and 15ns at intervals of 0.4ns for a repetition period of 25ns and for numbers of time-channels between 4 and 1024. We then convolved the exponential PDF with an IRF generated using the same instrument parameters and shaped as a Gaussian with a standard deviation at intervals between 0.01 and 1ns or as a rectangle of similar width. Finally, we added Poissonian noise to the resulting decay curves after rescaling to simulate a signal containing 75,000 photons. Unless otherwise described, the signal was then fit by iterative reconvolution using the simulated IRF and a Levenberg-Marquardt algorithm to obtain the lifetime estimate. We fitted the simulated data with the known IRF. However, to quantify the information losses caused by the lack of an experimental IRF, we also fit the width of the IRF for a subset of the simulations. We used the variance in the lifetime estimates of 5,000 simulations to evaluate F-values.

### Numerical and analytical estimates

Numerical and analytical methods used to characterise Fisher information were developed using Mathematica (Wolfram Research, Inc., Champaign, IL). These are freely available as Mathematica Notebooks at the GitHub repository *Esposito-Lab/BiochemicalResolution* [32]. The notebook ‘*DiracIRF_FisherInformation_V1.nb*’ introduces the formalism described already by the seminal papers of [6, 20]. The notebook ‘*GaussianIRF_FisherInformation_V1.nb*’ extends this formalism for the more general case of a Gaussian IRF of finite width. These files also include the code for the numerical evaluations of the F-value for the Gaussian IRF case study.

### System performance loss

We represented performance loss between two systems as the area between their F-value over lifetime curves. To evaluate the performance loss between Gaussian shaped IRFs and Dirac IRFs, we used the numerically derived F-values and evaluated over the lifetimes of 10ps and 15ns at intervals described by **Eq. (10)**. Due to numeric instabilities at lower lifetimes, differences between the Gaussian and Dirac IRFs that were negative were set to zero. To evaluate the performance loss between Gaussian and rectangular shaped IRFs and between measured and fitted IRFs, the Monte Carlo simulation derived F-values were used instead of the numerically derived F-values because of the challenges in obtaining analytical solutions for different IRF shapes and for fitted IRF conditions. Due to instabilities in simulating at faster decay constants, we evaluated over a shorter lifetime range of 0.6 to 15ns.

## 3. Results

### Fisher information theory

First, we have derived analytically the description of the Fisher information related to the amplitude and the fluorescence lifetime estimates for the case study of a generic TCSPC system with a Gaussian IRF. Briefly, we evaluated the convolution integral shown in **Eq. (2)** for a Gaussian with a standard deviation of *σ_irf_* centred at *t*_0_ and with an amplitude of the exponential of *A*.

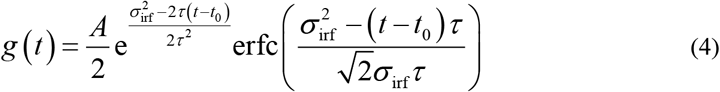

We then integrated **Eq. (4)** to evaluate the number of photons Λ (*t*) counted from an excitation pulse to time-delay *t*.

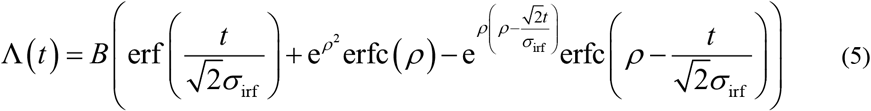

For the convenience of notation, we write **Eq. (5)** as a function of parameters *B* = *Aτ* / 2, 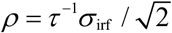 and *t*_0_ = 0. The analytical estimation of the photon counts detected in a given time-bin indexed by 1 ≤ *i* ≤ *m* is described by *G*(*i*) = Λ (*iT*) – Λ ((*i* – 1)*T*), where *G*(*i*) is used in our notation to indicate the expectation of Λ (*iT*) – Λ ((*i* – 1)*T*), and T is the period of the pulse train. With this notation, we can evaluate the log-likelihood analytically, as shown in **Eq. (6)**.

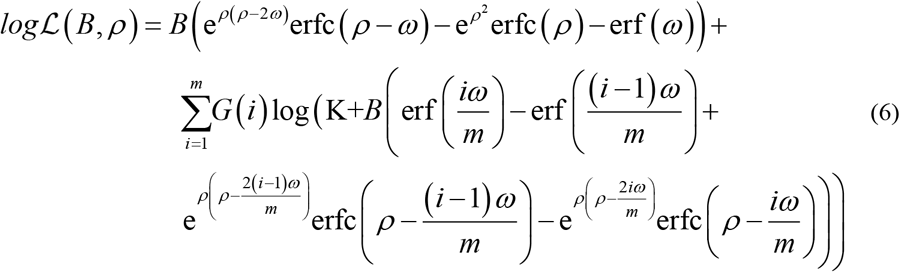

Here, we have introduced 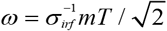 that, together with *B* and *ρ* permits a convenient parametrisation of the log-likelihood on a time base defined by the width of the IRF. We have also used a regularisation constant K, to avoid numerical instabilities when the argument of the logarithm might become 0. The Rao-Cramer [6, 24, 33] bound for an unbiased estimator of the fluorescence lifetime *τ* can be quantified with the study of the Fisher information matrix 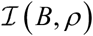, that in this case can be written as:

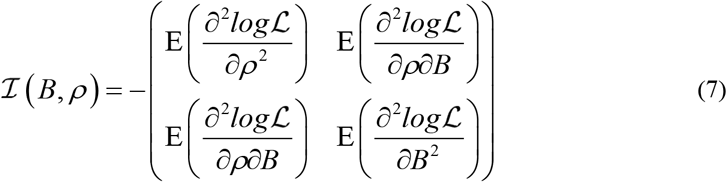

*E* is the expectation operator, which estimate for *G*(*i*) can be easily evaluated as in **Eq. (8)**.

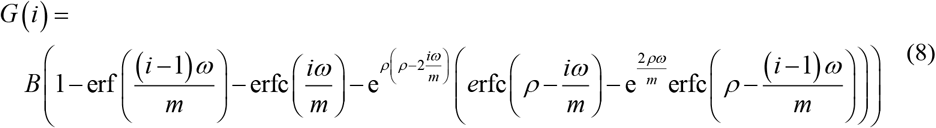

The top-left value of 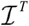 is the variance of *ρ*, which then permits to evaluate the F-value of a TCSPC system with a Gaussian IRF, substituting 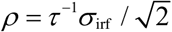 and 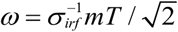.

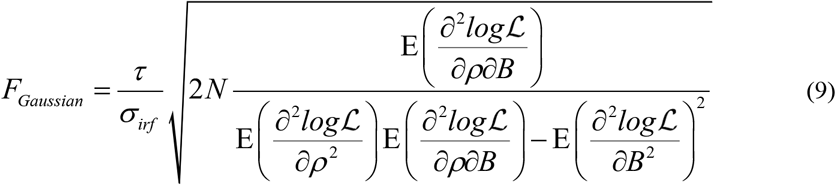

Here, we present numerical evaluations of *F_Gaussian_* (**Fig. 1(a-e)**) because we could not simplify the analytical solution (available in the supporting Mathematica Notebook) to an intuitive short form. The regularisation constant *K* was set to 10^-6^ not to alter the general solution and permitting divergence for low values of *τ* while avoiding indefinite forms. The solutions were evaluated numerically on a lattice of ~1.3 10^4^ parameter values defined by **Eqs. (10)**.

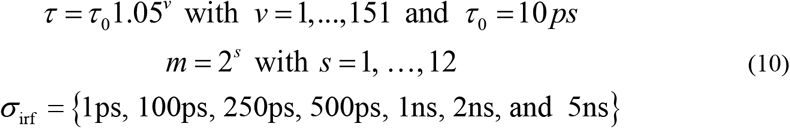

**Figure 1.**
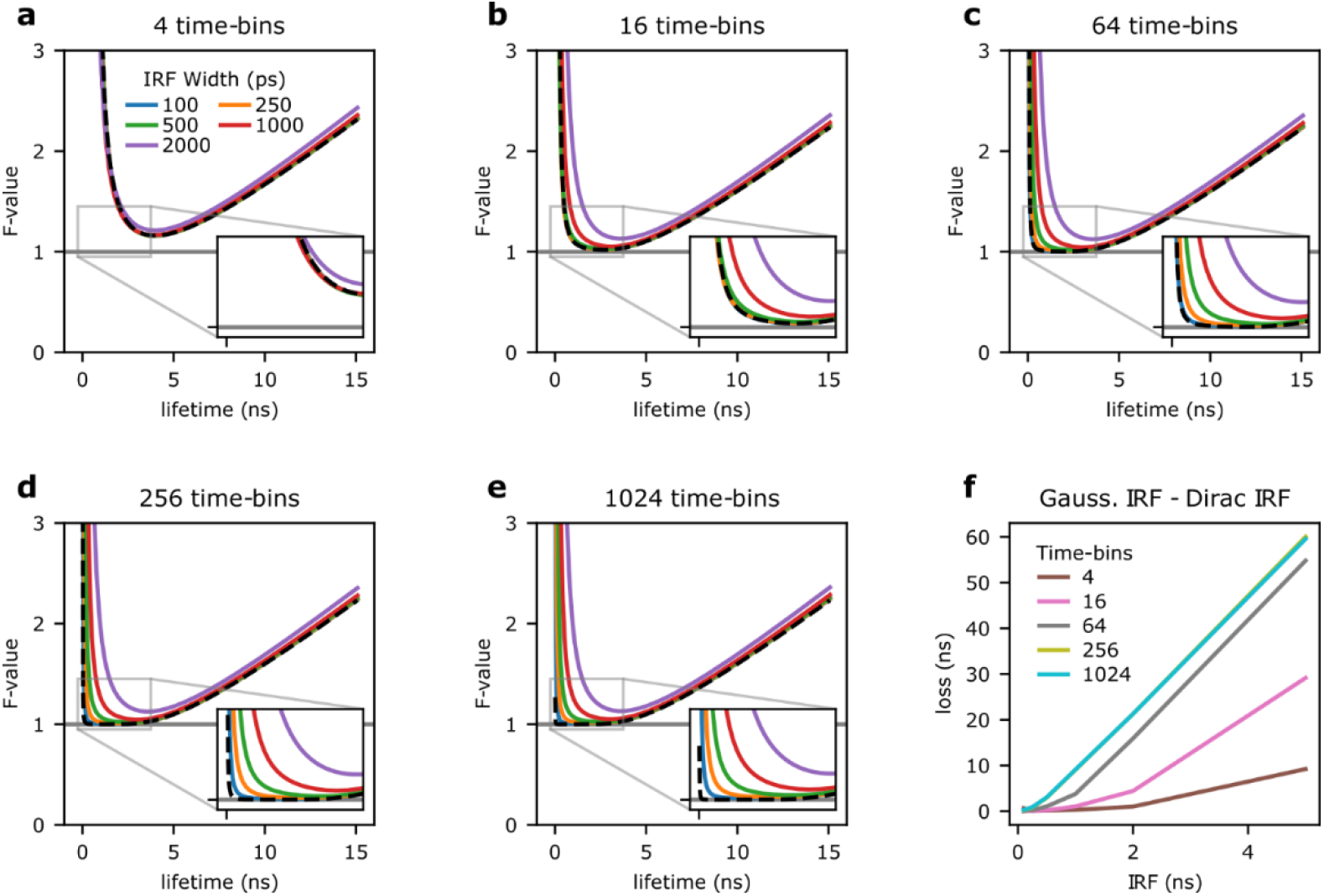
Fisher information and instrument response function. **a-e)** Fisher information analysis was used to estimate the photon economy (F-value) of TCSPC systems for Gaussian IRFs with standard deviations of 100ps (blue), 250ps (orange), 500ps (green), 1ns (red), and 2ns (purple). The F-value is equal to one when the signal is limited only by Poissonian noise (solid grey line). For reference, the F-value of a Dirac IRF for each system is also shown (black dashed line). The inserts show a magnified area of the plot (from −0.25 to 3.75ns, and for F-values between 0.95 and 1.45). Systems with 4 (a), 16 (b), 64 (c), 256 (d), and 1024 (e) time-bins for a 25ns measurement period were considered. **f)** The difference between the area under the curves shown in panels (a-e) for a Gaussian- and a Dirac-IRF is used as a measure of performance loss caused by non-ideal instrument response function.

In the rest of the work, the period of the pulse train is equal to 25ns.

### Increasing IRF widths reduce photon efficiency

**Fig. 1(a-e)** shows the F-value estimated with **Eq. (9)**, with the black dashed line representing the reference performance of a Dirac-excitation. For a time-resolved system with only four time-bins (**Fig. 1(a)**), similar to some time-gating implementations [12, 34], the F-values are nearly indistinguishable from the theoretical limits of a Dirac IRF (here *σ*_irf_ =100, 250, 500, 1000, and 2000ps are shown). For low time-bin resolutions (*Tm^-1^*), sampling limits performances, and the width of the IRF does not cause further losses in precision, at least until the IRF is comparable to the resolution of the photon-arrival times histogram (here *Tm^-1^* ~6ns/bin).

In **Fig. 1(b-e)**, the number of time-bins are increased to 16, 64, 256, and 1024 (equivalent to *Tm^-1^*~1.6ns, 0.4ns, 0.1ns, 0.02ns), respectively. As expected, the F-values for both the theoretical Dirac-limit and finite Gaussian IRFs improve as the number of time-bins increases, particularly at shorter decay times. However, the improvement for Gaussian IRFs with broader width (1-2ns) are more modest.

To further quantify the effects of the IRF, we evaluated the performance loss as the area under the curve between the Gaussian IRF and the theoretical Dirac-limit F-values. **Fig. 1(f)** again illustrates that while the performance loss generally increases with the IRF width, the loss is minimal until the IRF width is similar to the time-bin resolution. Therefore, for systems with fewer time-bins, a wider IRF can be tolerated before a loss in performance is observable. While the results shown in **Fig. 1** are somewhat expected, taken together, it illustrates that very narrow IRFs are only beneficial for very short fluorescence lifetime values and systems with a high time-bin resolution.It has been previously noted that an increase above 64 time-bins will only provide minor improvements in the photon economy of a system [20, 22]. We have then found that with a 64 time-bins system at 40MHz, an IRF up to 250ps can be tolerated with only minor deterioration in system performance. Increasing the number of time-bins beyond 64 and decreasing the IRF will not significantly improve the photon economy for most lifetimes over the measurement period; however, it may improve the photon economy at very short lifetimes. The capability of a TCSPC system to measure fluorescence lifetimes shorter than the IRF width has been reported in the past [35]. With these results, we can appreciate how this is both a function of instrument parameters and photon-budget. Short fluorescence lifetimes require an increasing number of photons to be characterised. Compared to the best performance achievable from a TCSPC (*F*=1), to achieve the same precision for a short lifetime, the detection of *F^2^* photons would be necessary. For shorter lifetimes, the loss of Fisher information due to an IRF wider than 250ps becomes so high that it makes any measurement impractical.

### The shape of the IRF determine TCSPC photon-efficiency

Next, we cross-validated the numerical estimations of **Eq. (9)** (**Fig. 1(a-c)**) with Monte Carlo simulations (**Fig. 2**; 4, 16, and 64 time-bins) achieving a very close match between the two independent methods. Having shown the excellent performance of Monte Carlo simulations, we used this approach to understand the dependencies of TCSPC photon-efficiencies on different parameters because analytical solutions for general cases can be arduous to obtain. In **Fig. 3(a-c)**, we compare the F-value of Gaussian- (solid curves), and rectangular- (dashed curves) shaped IRFs of equal full-width at half-maximum in 16 and 256 time-bin systems. For a Gaussian, the FWHM is equal to 2.355σ [36]. For narrow FWHM values (*e.g*., 100ps), the F-value of both Gaussian and rectangular cases are very similar converging, unsurprisingly, to the Dirac solution. However, with the increasing width of the IRF, the photon-efficiency of the Gaussian-shaped IRF deteriorates faster than the rectangular IRF as shown in **Fig. 3(d)**. In comparing 16 to 256 time-bin systems, higher time-bin resolution decreases the system performance loss due to IRF shape. The comparison of performance between Gaussian and rectangular IRF of equal FWHM leads to two conclusions. Trivially, we observe that the shape of the IRF impacts the photon-efficiency of a TCPSC system significantly. More interesting, rectangular IRFs of low but finite duty-cycle can outperform systems with Gaussian IRFs.

**Figure 2.**
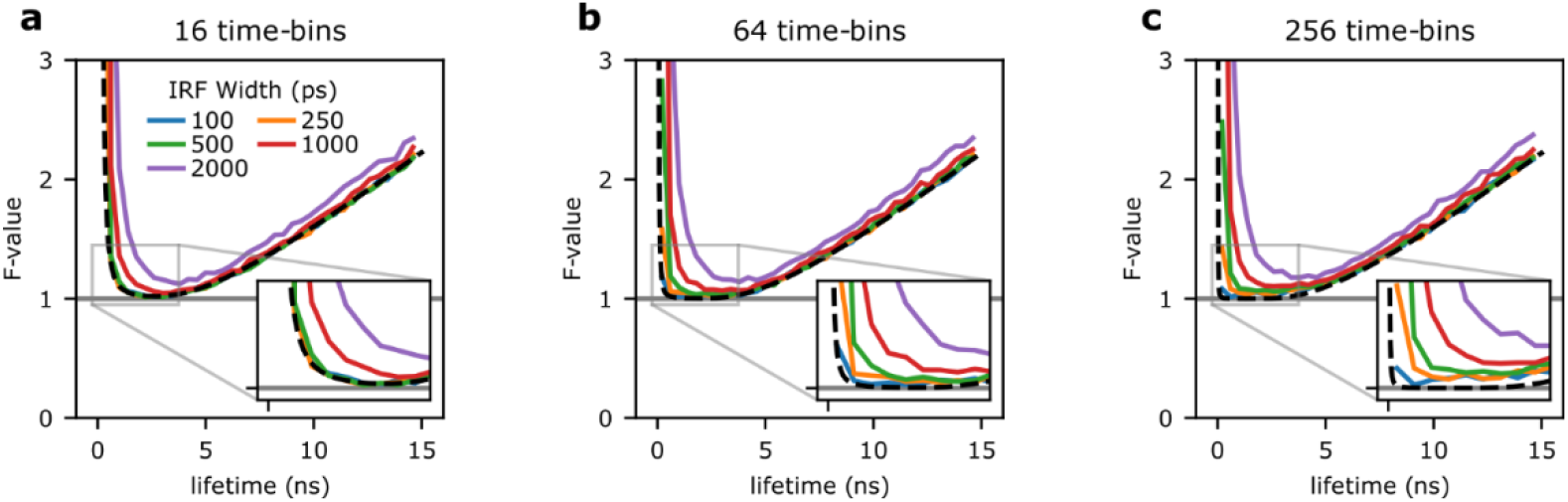
The impact of the IRF on photon economy studied by Monte Carlo simulations. **a-c)** Monte Carlo simulations were performed to cross-validate the Fisher information analysis shown in **Fig. 1**. Fluorescence lifetime signals were simulated and lifetimes were estimated for systems with Gaussian IRFs with standard deviations of 100ps (blue), 250ps (orange), 500ps (green), 1ns (red), and 2ns (purple). Each simulation run comprised 75,000 photon-events and was repeated 5,000 to calculate F-values. Systems with 16 (a), 64 (b), and 256 (c) time-bins sampling a 25ns measurement period were considered. The theoretical F-value, of a Dirac IRF for each system is shown as reference (black dashed line).

**Figure 3.**
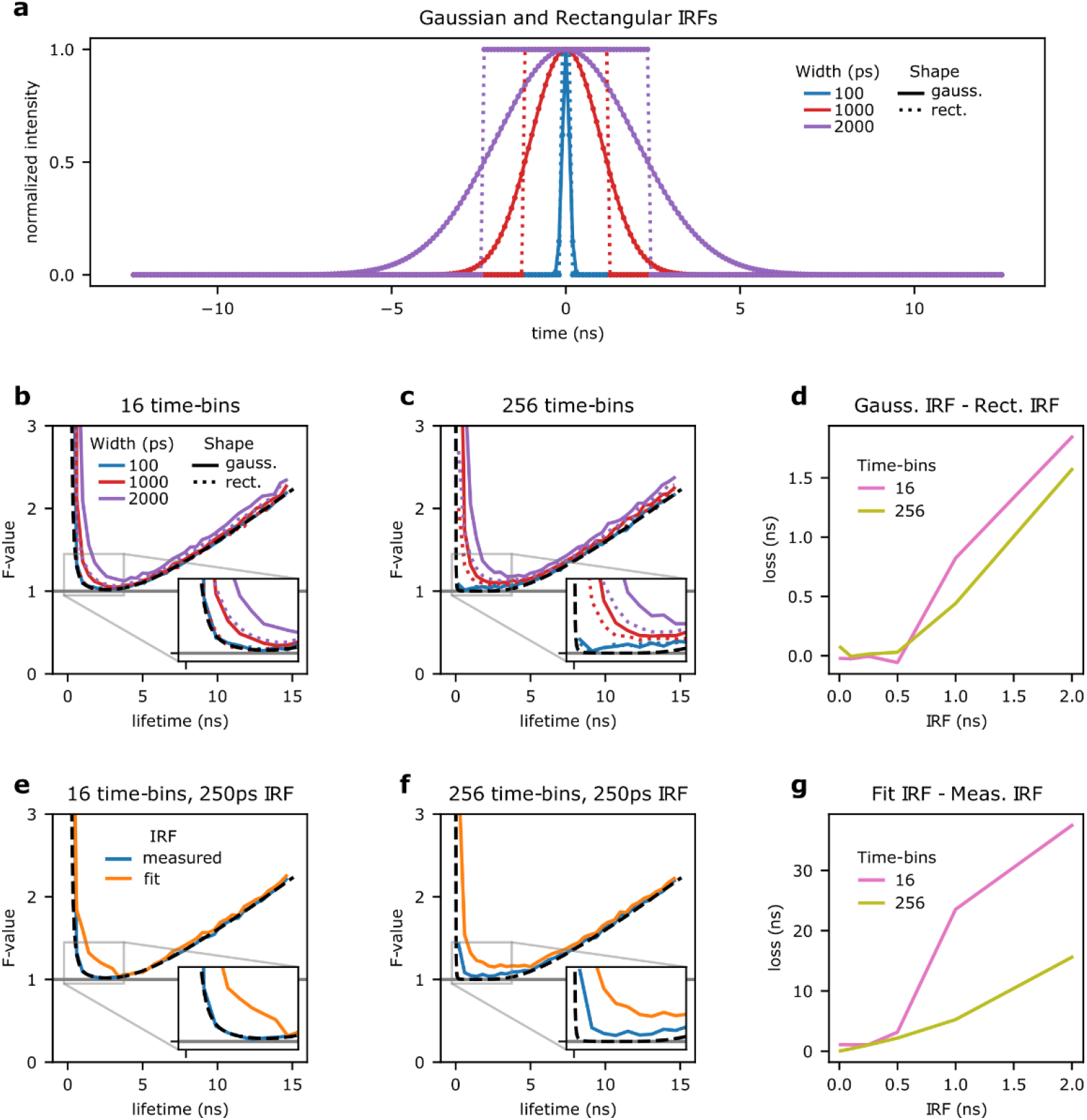
The impact of the IRF shape and measurement on photon economy. **a-c)** Monte Carlo simulations were performed to compare the photon economy between Gaussian and rectangular shaped IRFs. Gaussian IRFs (solid lines) with a standard deviation of 100ps (blue), 1ns (red), and 2ns (purple) were compared to rectangular IRFs (dashed lines) at equivalent FWHM. These IRFs are shown for a system of 256 time-bins (a). The photon economy was compared in systems of 16 (b) and 256 (c) time-bins. **d)** The differences between the areas under the curves represent the performance loss for Gaussian-relative to rectangular-instrument response functions for both systems with 16 (pink) and 256 (yellow-green) time-bins. **e-f)** Monte Carlo simulations were also performed to compare the photon economy of iterative reconvolution data fitting where the IRF width is estimated by fitting (orange) or experimentally (blue). For example, the measurement is simulated with a Gaussian IRF with a standard deviation of 250ps in a system with 16 (e), and 256 (f) time-bins. **g)** The performance loss caused by fitting the IRF width is shown for both systems with 16 (pink) and 256 (yellow-green) time-bins.

We have shown that an IRF below 250ps does not have a significant impact on fluorescence lifetime estimates. This observation is valid when the IRF is well-known *a priori* and used in iterative reconvolution fitting [17, 18, 37]. However, fitting the IRF width causes a considerable loss of precision of fluorescence lifetime estimates for lifetime values of any magnitude. For example, **Fig. 3(e-f)** illustrates the information cost of estimating the IRF width rather than measuring it for a Gaussian IRF of 250ps in 16 and 256 time-bin systems. Similar to the case with IRF shape, higher time-bin resolution decreases the system performance loss due to estimating relative to measuring the IRF as shown in **Fig. 3(g)**. The experimental characterisation of the IRF is thus essential not just to improve the accuracy of fluorescence lifetime estimates (the fit of a wrong IRF model results in inaccurate decay values) or the minimal fluorescence lifetime that can be resolved but also the precision of fluorescence lifetime estimates more generally.

### The impact of the IRF onto the biochemical resolving power of a microscope

Fluorescence lifetime imaging microscopy is often used in biomedical research to probe cellular biochemistry. In biochemical imaging applications, the measurement of absolute values of fluorescence lifetimes is often of little interest. Instead, it is often the ability to distinguish between spatiotemporal variations of a fluorescence lifetime estimate (*e.g*., the comparison between two pixels or the time response within a pixel) that is more relevant. Köllner and Wolfrum defined separability (*S*) as a measure of the capability of a system to differentiate between two lifetime measurements, τ_1_ and τ_2_: [20]:

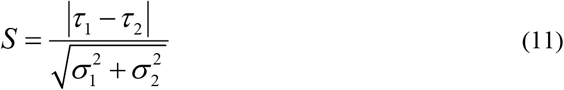

where σ_1_ and σ_2_ are the standard deviations of the measurements of τ_1_ and τ_2_, respectively. To illustrate how separability is affected by the IRF, we used our numerical estimations of F-values to express separability by assuming the collection of 2,000 photons per measurement. Using **Eq. (11)**, we compared the separability of two hypothetical probes with a dynamic range of 10%; the first probe operates in the lower lifetime region with a range between 0.50 and 0.55ns and a second probe in the higher lifetime region with a range between 2.0 and 2.2ns (**Fig. 4(a)**).

**Figure 4.**
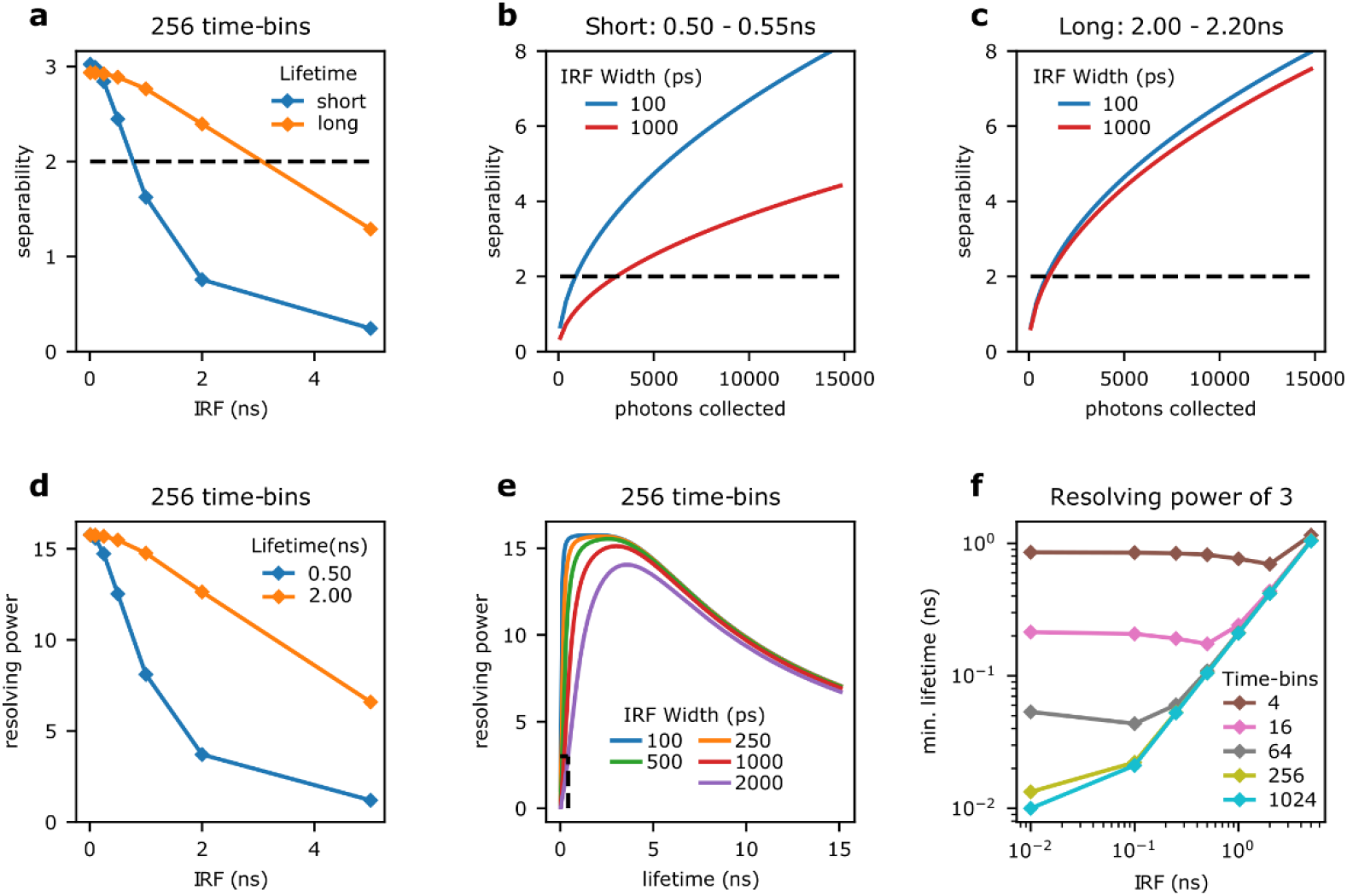
The biochemical resolution of fluorescence lifetime imaging. **a)** The separability of two fluorescence lifetime decays decreases for broad instrument response functions, here shown for a photon-budget of 2,000 photons collected with a system of 256 time-bins. We show instances of two probes exhibiting fluorescence lifetime variations of 10% for a sensor exhibiting a short fluorescence lifetime (switching from 0.50 to 0.55ns; blue curve) and a longer one (from 2.0 to 2.2ns; orange curve). **b-c)** The increase in collected photons needed to maintain separability between a Gaussian IRF with a standard deviation of 100ps (blue) and 1ns (red) was evaluated for both probes. In panels a-c) we show the reference separability of 2 (dashed black line) where two fluorescence lifetimes can be considered distinguishable. **d)** The resolving power of a fluorescence lifetime imaging system is shown for reference fluorescence lifetimes of 500ps (blue) and 2ns (orange) as a function of IRF width, evaluated for a photon-budget of 2,000 photons collected with a system with 256 time-bins. **e)** From plots of resolving power as function of fluorescence lifetime and IRF width (100ps (blue), 250ps (orange), 500ps (green), 1ns (red), and 2ns (purple)), we inferred the smallest resolvable fluorescence lifetime to achieve at least a resolving power of 3 **(f)**. The dashed curves shown in panel e) show the reference line of R=3 (horizontal line) and the correspondent resolution limit (vertical line) for a broad IRF of 2ns width (purple).

For narrow Dirac-like IRFs, the separability between the two probes are similar due to both distinguishing a 10% change in lifetimes. As the IRF width increases, the separability of both probes decreases. However, the probe operating at longer lifetimes is less affected by the increasing IRF. The longer lifetime probe remains above a separability of 2 up to an IRF width of approximately 2.0ns, while the shorter lifetime probe can only tolerate up to an IRF of approximately 0.5ns.

To increase the separability, more photons can be collected per measurement, as shown in **Fig. 4(b-c)**. The increase in separability as additional photons are collected is shown for the probe operating in the lower lifetime region (**Fig. 4(b)**). For this probe, increasing the IRF width from 0.1 to 1.0ns requires an increase from approximately 1,000 to 10,000 photons to maintain the same level of separability, which translates to the same fold increase in image acquisition time. In comparison, for the probe operating in the higher lifetime range, the same increase in the IRF requires a significantly smaller increase in photons collected to maintain the separability (**Fig. 4(c)**).

While separability quantifies the ability to distinguish between two fluorescence lifetime estimates, the biochemical resolving power describes the smallest change (Δ*τ*) that can be measured for a given fluorescence lifetime *τ*_0_. We have previously defined the biochemical resolving power as *R* = *τ*_0_ / Δ*τ*, which is equivalent the definition of resolving power in spectroscopy, *R* = *λ* / Δ*λ* [24]. In analogy to the spatial resolution of a microscope, we have previously derived the biochemical resolving power using the Rayleigh criterion, which corresponds to a separability of 2. By substituting *τ_1_*, *τ_2_*, and *σ_1_*=*σ_2_*=*σ_0_* with *τ_0_*+*Δτ/2*, *τ_0_-Δτ/2*, and 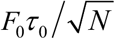, respectively, the biochemical resolving power is expressed as (see Ref [24] for the full derivation):

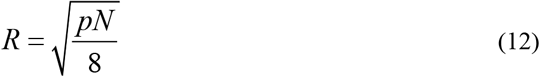

where the photon-efficiency of a FLIM system, *p* = *F*^-2^ [23], and the number of detected photons, *N*, determine the biochemical resolving power of a microscope [24].

Using **Eq. (12),** we compared the effects of an increasing IRF on resolving power between a lifetime of 0.5 and 2.0ns. **Fig. 4(d)** shows that the resolving power decreases at a faster rate for a shorter lifetime. **Fig. 4(e)** shows the dependency of *R* on both fluorescence lifetime and IRF width. We can use this parametric representation to infer which is the minimum measurable fluorescence lifetime as a function of IRF width, number of bins and photon-counts. For example, in **Fig. 4(e)** we show the resolution limit (vertical dashed lines) achievable with a target *R* value set to 3 (horizontal dashed lines). **Fig. 4(f)** shows the smallest measurable fluorescence lifetime values with *R*=3 for different sampling resolutions and instrument response function widths. These minimum lifetimes are also summarized in **Table 1**. Replacing the photon-efficiency for a given lifetime in **Eq. (12)** and solving for *N* (*i.e*., *N* = 8*p*^-1^*R*^2^) we can then estimate the number of photons necessary to reach this resolution limit. For example, a TCSPC system with a Gaussian IRF of 100ps, 256 time-bins and a laser repetition period of 25ns could resolve a fluorescence lifetime of ~20ps with a target *R* of 3. In these conditions *p* is ~0.03; thus, the number of photons necessary to resolve a short fluorescence lifetime of 20ps would be ~2,700. Other combinations of minimum resolvable lifetimes and IRF widths are summarized in **Table 2**.

**Table 1.** Minimum Resolvable Lifetimes for Time-bin and IRF Combinations. Minimum resolvable lifetimes. The smallest resolvable fluorescence lifetimes achievable with a resolving power of 3 and a photon budget of 2,000 were evaluated for various time-bin and IRF combinations.

|  | Time-bins |  |  |  |  |
| --- | --- | --- | --- | --- | --- |
|  | 4 | 16 | 64 | 256 | 1024 |
| IRF (ps) | Minimum Lifetime (ns) |  |  |  |  |
| 100 | 0.85 | 0.21 | 0.04 | 0.02 | 0.02 |
| 250 | 0.84 | 0.19 | 0.06 | 0.05 | 0.05 |
| 500 | 0.82 | 0.17 | 0.11 | 0.11 | 0.10 |
| 1000 | 0.76 | 0.24 | 0.21 | 0.21 | 0.21 |
| 2000 | 0.70 | 0.44 | 0.42 | 0.42 | 0.42 |

**Table 2.** Required Photons for Minimum Resolvable Lifetimes for IRF Widths. Photons required for minimum resolvable lifetimes. The number of photons required were evaluated for a given minimum resolvable fluorescence lifetimes and IRF width that is achievable with a resolving power of 3.

|  | Minimum Resolvable Lifetime (ns) |  |  |  |  |
| --- | --- | --- | --- | --- | --- |
|  | 0.02 | 0.05 | 0.10 | 0.25 | 0.50 |
| IRF (ps) | Photons Collected |  |  |  |  |
| 100 | 2,710 | 290 | 120 | 80 | 70 |
| 250 | 22,570 | 2,320 | 430 | 110 | 80 |
| 500 | 104,470 | 12,680 | 2,330 | 270 | 110 |
| 1000 | >1,000,000 | 61,800 | 12,890 | 1,280 | 270 |
| 2000 | >1,000,000 | >1,000,000 | 62,470 | 7,290 | 1,310 |

Taken together, the data shown in **Fig. 4** characterises the two fundamental aspects of ‘resolution’ for time-resolved instrumentation: i) the ability to separate two different fluorescence lifetimes and ii) the smallest measurable fluorescence lifetime. While the latter definition (ii) is commonly used as a reference for resolution, oftentimes in biology, the ability to separate two lifetimes is what matters. Both properties are affected by specification of instruments, but we show that the ability to separate lifetimes in the range that are often used for biological applications are more lenient on the required IRF and time-bin requirements.

## 4. Discussion

The role of photon-statistics for time-resolved measurements has been widely characterised. However, to our knowledge, a quantitative description of how the instrument response function contributes to the precision of a measurement and ultimately limits the resolution of a time-resolved technique was lacking. To fill this knowledge gap, we have applied Fisher information theory and Monte Carlo simulations to evaluate the role of IRFs on photon economy. First, we described the theoretical considerations which characterised the performance of a time-resolved system from the perspective of decay-time resolution which offers provocative practical considerations.

Unsurprisingly, a narrower IRF and denser sampling of the fluorescence decays improve the overall system performance converging to the ideal performance (*F*=1). However, for a given photon-budget, the improvements achievable by narrower IRFs are limited by the histogramming bin resolution. Notably, for typical high-end time-tagging electronics, timing precisions and the minimum time-bin widths are positively correlated. However, the impact on the photon-efficiency of a time-resolved system also depends on the dead-time of the electronics and detectors (higher dead-times, higher photon-losses), the transit time spread of the detectors (typically in the range of 20-200ps) which will contribute to the system IRF, and the quantum efficiency of the detector. For example, when seeking to maximise the photon-budget by selecting detectors with high quantum efficiencies and electronics with low dead-times, compromises might be necessary on the nominal system IRF.

Leading manufacturers of TCPSC systems supply hybrid photo-multiplier tubes (PMTs) such as the bialkali HPM-100-06 (Becker-Hickl GmbH) and PMA-06 (PicoQuant GmbH) that exhibit excellent dead-times (<1ns), very low transit time spread (20-50ps), and good quantum efficiency spanning the UV and visible spectrum (a peak 28% at 400nm). The equivalent packaged detector with a GaAsP photocathode (HPM-100-40 or PMA-40) significantly improves the quantum yield of such devices in the visible range (from <20% for the bialkali to ~45% for the GaAsP). With all other experimental parameters equal, using a GaAsP photocathode instead of a hybrid PMT could provide up to three times higher photon-counts for a green fluorophore. From **Eq. (3)**, we can then infer that for a shot-noise limited system, we could achieve a 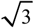 improvement in the (biochemical/lifetime) resolving power of a detection system. However, the −40 series of hybrid PMTs exhibit a much higher (120ps) transit time spread compared to the −06 series. Our work would then suggest that for nanosecond-lived fluorescence lifetimes (or for lifetimes >500ps in this case), the loss in resolving power caused by the IRF is insignificant and the 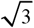 improvement determines the performance of the system. However, when measuring shorter lifetimes (<50ps), assuming that the time-tagging electronics provide sufficient time resolution (>256 time-bins), the transit time spread of the GaAsP photo-cathode will incur a high enough penalty that it cannot be offset by the 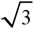 improvement in collection efficiency.

Far from the limit of resolution (the smallest decay value that could be measured), which depends in non-straightforward ways on the various parameters discussed, the biochemical resolving power of a FLIM system is affected more dramatically from the prior knowledge of the IRF rather than it being extremely narrow. Indeed, we have shown how broader IRFs and even rectangular-shaped IRFs could provide resolving powers as high as ideal Dirac-like excitation profiles. This observation is reminiscent of a previous observation reported for frequency-domain fluorescence lifetime microscopy [23], where rectangular excitation pulses of finite width (1-2ns) could provide higher performances compared to typical sinusoidal excitation, permitting a system to approach the ideal performances of a time-domain FLIM with Dirac excitation. Similarly, we can now infer that for a system in which the IRF is dominated by the width of the excitation pulse, broad nanosecond pulses might not significantly deteriorate the performance of a FLIM system provided that the IRF is experimentally well-characterised, stable and the fluorescence lifetime decays are sampled sufficiently well.

We note that when fitting a wrong IRF model (for example using a Gaussian approximation for non-Gaussian IRFs) the accuracy of lifetime estimates deteriorates. However, while systematic errors might be tolerated in applications where only changes in fluorescence lifetimes are relevant, the loss of precision and resolution caused by estimating IRF parameters by fitting on a pixel-by-pixel basis is vastly detrimental. Although an analysis of accuracy is beyond the scope of this work, we note that the experimental characterisation of the IRF is thus always advisable not only to retrieve the most accurate results but also to maximise the precision and resolution of an assay.

Some of these observations might be considered intuitive, our results have both practical implications (*e.g*., as described for the selection of a photo-cathode material) and provocative conclusions.

First, ambiguity between different definitions or types of resolutions should be avoided. For all practical purposes, in the biomedical field, it is what we define as biochemical resolving power of a microscope that is relevant. This is a quantity that is limited by the photon-budget and for experiments using typical organic, inorganic (*e.g*., Quantum Dots), or genetically encoded fluorophores and high-end TCSPC systems, the IRF width and shape might have a smaller impact onto the biochemical resolving power of an instrument than often perceived.

Second, for very short-lived fluorophores, the impact of the IRF is naturally significant. However, the biochemical resolving power is still photon-budget limited whenever the minimum bin width is sufficient for sampling the IRF and the fluorescence decay. In this case, the limit of resolution might be better expressed in terms of the smallest value that could be measured with a given photon-budget. In fact, the capability for a TCSPC to quantify fluorescence decays with time constants shorter than the width of the IRF has been illustrated experimentally in the past with the collection of approximately 1,000 photons in the maximum time channel [35, 38].

Third, maximum care should be taken in the collection of as much signal as possible from the limited photons that typical fluorophores used in biomedical application can emit. As the photon-budget will ultimately determine the biochemical resolving power in typical assays (*e.g*., sensors based on fluorescent proteins), the excellent resolving power that could be provided by the high timing-resolution of high-end TCSPC electronics and femtosecond lasers can be squandered by the use of less sensitive detectors or optical relay systems.

We also note that time-resolved systems are not only utilised in biomedical applications or fluorescence. Phosphorescence lifetime imaging is often applied to sense molecular oxygen in tissues using much longer lifetimes than fluorescence [39]. In comparison, extremely short delay times are measured in a large variety of instrumentations spanning nuclear physics to mass spectrometers, and also include 3D ranging technologies used for machine vision and in the automotive industry. As the theory that we have described here can be directly applicable or can be adapted to the description of resolving power of timing of any of these applications, we envisage that this work might expand our understanding and could be used for the improvement of several time-resolved technologies across different industries.

## Acknowledgments

AE acknowledges the financial support provided by the CRUK with a multi-disciplinary project award (OncoLive, C54674/A27487). AE and ALT acknowledges financial support from Medical Research Council program grants (MC_UU_12022/1 and MC_UU_12022/8) awarded to Prof. Ashok Venkitaraman.

